# Class A Penicillin-Binding Protein-mediated cell wall synthesis promotes structural integrity during peptidoglycan endopeptidase insufficiency

**DOI:** 10.1101/2020.07.03.187153

**Authors:** Shannon G. Murphy, Andrew N. Murtha, Ziyi Zhao, Laura Alvarez, Peter Diebold, Jung-Ho Shin, Michael S. VanNieuwenhze, Felipe Cava, Tobias Dörr

**Author notes:** Department of Biochemistry and Biophysics, University of California, San Francisco, San Francisco, CA 94158, USA.

## Abstract

The bacterial cell wall is composed primarily of peptidoglycan (PG), a poly-aminosugar that is essential to sustain cell shape, growth and structural integrity. PG is synthesized by two different types of synthase complexes (class A Penicillin-binding Proteins [PBP]s/Lpos and Shape, Elongation, Division, Sporulation [SEDS]/class B PBP pairs) and degraded by ‘autolytic’ enzymes to accommodate growth processes. It is thought that autolsyin activity (and particulary the activity of endopeptidases, EPs) is required for PG synthesis and incorporation by creating gaps that are patched and paved by PG synthases, but the exact relationship between autolysins and the separate synthesis machineries remains incompletely understood. Here, we have probed the consequences of EP depletion for PG synthesis in the diarrheal pathogen *Vibrio cholerae*. We found that EP depletion resulted in severe morphological defects, increased cell mass, a decline in viability, and continuing (yet aberrant) incorporation of cell wall material. Mass increase and cell wall incorporation proceeded in the presence of Rod system inhibitors, but was abolished upon inhibition of aPBPs. However, the Rod system remained functional (i.e., exhibited sustained directed motion) even after prolonged EP depletion, without effectively promoting cell elongation. Lastly, heterologous expression of an EP from *Neisseria gonorrhoeae* could fully complement growth and morphology of an EP-insufficient *V. cholerae*. Overall, our findings suggest that in *V. cholerae*, the Rod system requires endopeptidase activity (but not necessarily direct interaction with EPs) to promote cell expansion and substantial PG incorporation, whereas aPBPs are able to engage in sacculus construction even during severe EP insufficiency.

**Importance:** Synthesis and turnover of the bacterial cell wall must be tightly co-ordinated to avoid structural integrity failure and cell death. Details of this coordination are poorly understood, particularly if and how cell wall turnover enzymes are required for the activity of the different cell wall synthesis machines. Our results suggest that in *Vibrio cholerae*, one class of turnover enzymes, the endopeptidases, are required only for substantial PG incorporation mediated by the Rod system, while the aPBPs maintain structural integrity during endopeptidase insufficiency. Our results suggest that aPBPs are more versatile than the Rod system in their ability to recognize cell wall gaps formed by autolysins other than the major endopeptidases, adding to our understanding of the co-ordination between autolysins and cell wall synthases. A detailed understanding of autolysin biology may promote the development of antibiotics that target these essential turnover processes.

## Introduction

Most bacteria elaborate a cell wall composed primarily of peptidoglycan (PG), which consists of polymerized N-acetyl glucosamine-N-acetyl muramic acid (poly-GlcNAc-MurNAc) dimers. These polymerized strands are covalently linked to each other via their oligopeptide side stems extending from the MurNAc residues; the degree of crosslinking varies with bacterial species and growth conditions (1–3). As such, PG encases the cell in a net-like structure that functions to maintain the high intracellular pressure accumulating in most bacteria and thus to prevent the cell from lysing. In concert with maintenance of structural integrity, PG has to accommodate growth processes (cell elongation and size expansion) and is therefore constantly degraded and resynthesized (4–6).

In many rod-shaped Gram-negative bacteria, cell wall synthesis during cell elongation is mediated by two separate types of cell wall synthase complexes: the Rod complex (which includes the glycosyltransferase RodA in conjunction with a class B Penicillin-Binding Protein [bPBP] and accessory proteins) and the class A PBPs in conjunction with their lipoprotein activators (7–12). The differential physiological roles of these PG synthases have only recently been begun to be dissected (13, 14), but remain incompletely understood.

Cell wall degradation, on the other hand, is mediated by a plethora of so-called “autolysins”, *i.e*., enzymes with the capability to break bonds in the PG sacculus (15–17). Members of one such group of autolysins, the endopeptidases (EPs), cleave the oligopeptide crosslinks between PG strands, presumably to allow for insertion of new PG material during cell elongation (18–21). To ensure structural integrity, EP-mediated cell wall cleavage and Rod- and/or aPBP-mediated resynthesis should logically be tightly coordinated, and this has indeed been demonstrated for cell elongation in Gram-positive bacteria (22–25). When the putative coordination is perturbed (*e.g*., after exposure to a cell wall synthesis inhibitor), PG structural integrity often catastrophically fails and cells die (26); this is one of the reasons why cell wall synthesis inhibitors (e.g., the β-lactams) rank highly among our most powerful antimicrobials (27). EPs in particular are a double-edged sword as they can both promote cell wall synthesis (28) and play major roles in cell wall cleavage after beta lactam exposure (29, 30). However, how EPs are regulated has only begun to be unravelled (31–34), and, at least in Gram-negative bacteria, we lack a complete understanding of how EP cleavage activity relates to PG synthesis by the two distinct cell wall synthase complexes.

Several models have been advanced to explain coordination of synthesis and degradation, with a prominent model being a “make before break” mechanism, where a nascent PG layer scaffold is elaborated parallel to an existing one, followed by cleavage of the old material that has been relieved of its critical structural function through this nascent PG load-bearing stabilizer (35, 36). Alternatively, PG might be able to sustain several cleavage events without experiencing catastrophic structural failure, obviating the need for any coordination between synthesis and degradation for as long as the Rod system and/or aPBPs are efficient enough in recognizing gaps in PG, *e.g*., through interaction with their cognate OM-localized activators in case of the aPBPs (a “break before make” model).

Here, we show that in the cholera pathogen *Vibrio cholerae*, EP activity is not required for cell wall synthesis *per se*. During EP insufficiency, growth and PG accumulation continue in the presence of Rod system inhibitors but abruptly stop upon inhibition of aPBPs. However, the Rod system continues directed motion for extended periods of EP depletion. Lastly, a heterologously expressed EP can fully complement growth and morphology of an EP-deficient *V. cholerae* strain. Our data suggest that aPBPs do not require wild-type levels of crosslink cleavage for PG incorporation (and consequently cell expansion), and that EP activity is required for the Rod system to contribute substantially to proper rod-shaped growth. Our cross-species complementation experiments intriguingly raise the possibility that direct co-ordination between EPs and cell wall synthases might not be necessary at all, at least under standard laboratory growth conditions.

## Results

### *Cell wall incorporation and mass increase continue during endopeptidase insufficiency in* V. cholerae

Endopeptidase depletion was previously shown to preclude insertion of new cell wall material in *E. coli*, resulting in rapid cell lysis (21). In contrast, we have noticed during EP depletion experiments with *Vibrio cholerae* that the cholera pathogen did not lyse, even in the absence of all 6 of its major D,D-EPs. This Δ6 endo strain (Δ*shyA* Δ*shyB* Δ*shyC* Δ*vc1537* Δ*vc0843*, Δ*vca1043* P_IPTG_:*shyA*), has the remaining, conditionally essential EP ShyA under control of an IPTG-inducible promoter and is thus suitable for depletion experiments. Upon growing the Δ6 endo strain in the absence of inducer (reducing ShyA to less than 10 % of initial levels after ~ 2h, **Fig. S1**), mass increase (measured by OD_600_) continued at a rate similar to when *shyA* expression was induced by IPTG (**Fig. 1A**). When plated on solid medium containing inducer at various timepoints after initiation of depletion, however, we observed a slight decrease in cfu/mL (**Fig. S2A**). Thus, the ability to recover and form colonies on a plate decreases during EP insufficiency. Additionally deleting the genes encoding PBP4, PBP7 and VC1269 (which have predicted EP activity but are, based on *E. coli*, not required for growth and cell elongation (37, 38)) did not appreciably affect mass increase (except for a slight decrease in final yield after 6 h; for an unknown reason, the strain also exhibitied a more pronounced drop in cfu/mL than Δ6 endo) (**Fig. S2B**), demonstrating that the mass increase phenotype did not simply reflect the ability of these putative EPs to substitute for ShyA.

**Figure 1.**
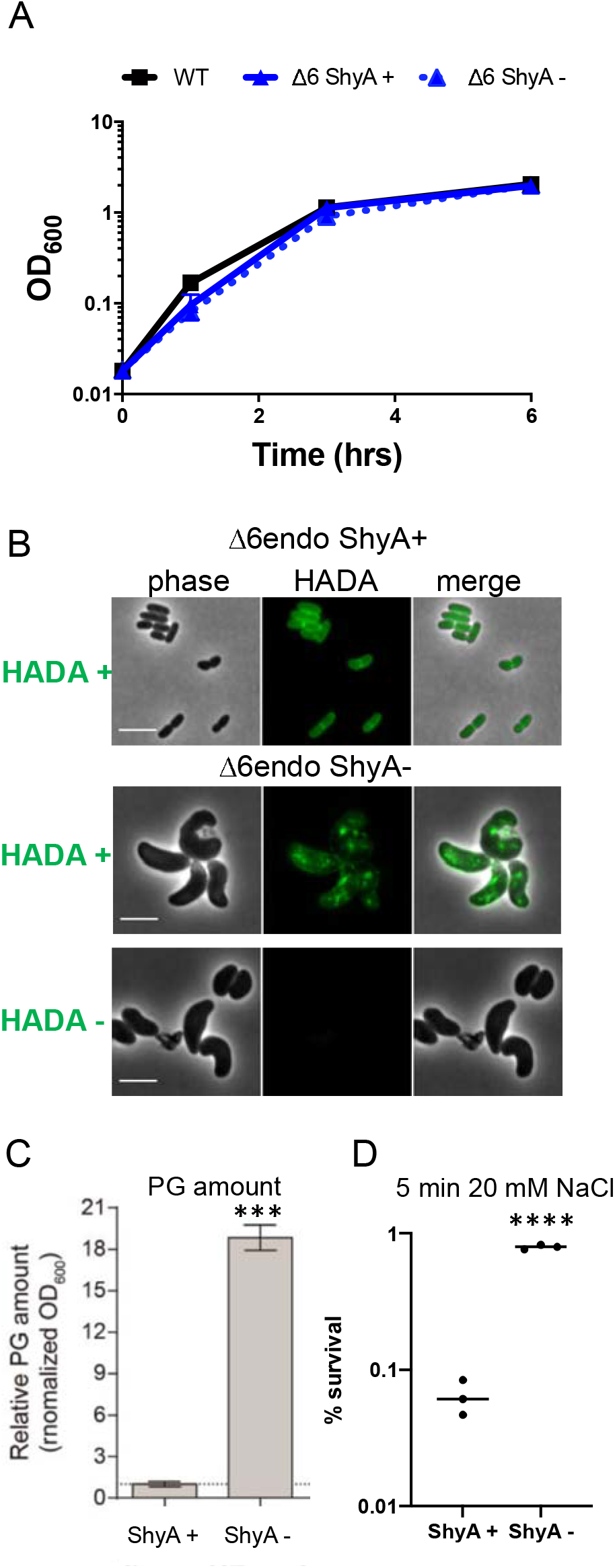
Cell mass increase and cell wall incorporation continue during EP insufficiency. **(A)** Overnight cultures of Δ6 endo (Δ*shyABC* Δ*vc1537* Δ*tagE1,2* P_IPTG_:*shyA*) grown in the presence of IPTG (200 μM) were washed twice and diluted 100-fold into growth medium with (ShyA +) or without (ShyA -) IPTG. At the indicated time points, OD600 was measured. Data are mean of six biological replicates, error bars represent standard deviation. **(B)** Δ6 endo was treated as described in (A) in the presence of HADA (100 μM). After 3 h, cells were washed twice and then imaged. **(C)** Relative PG content of Δ6 endo was measured via UPLC analysis (see methods for details) after 2 hours of growth in the presence (ShyA +) or absence (ShyA -) of inducer. Error bars represent standard deviation of 3 biological replicates. **(D)** Cells were treated as described in (A). After 3 h of growth in the presence or absence of inducer, cells were pelleted and resuspended in 20 mM NaCl (osmotic shock treatment) for 5 min. Shock treatment was stopped by adding PBS to 180 mM. % survival is cfu/mL before treatment divided by cfu/mL after treatment. Raw data points of 3 biological replicates are shown. **(C-D)** Asterisks denote statistical difference via unpaired t-test (****, p < 0.0001; ***, p < 0.001, ** p < 0.01, * < 0.05). Scale bars, 5 μm

We have previously shown that EP depletion in the Δ6 endo strain results in a dramatic increase in cell diameter and ultimately the generation of giant, bulky and contorted cells (34). Here, we probed the impact of endopeptidase insufficiency on PG composition and incorporation. PG architecture analysis revealed, as expected, that Δ6 accumulated a higher percentage of PG crosslinks after ShyA depletion (38.5%) compared to when ShyA is present (29.3 %), presumably due to the lack of EP cleavage activity (**Fig. S2C-D**). The analysis further revealed a 65% increase in trimer formation, as well as a 32% increase in the amount of anhydro residues upon ShyA depletion (**Fig. S2C-D**). Overall, the increase in crosslinking provides additional evidence that functional EP availability was highly limited under our depletion conditions. We next asked whether these enlarged cells elaborated a wild-type PG cell wall. To probe this, we cultured Δ6 endo in the presence of a fluorescent D-amino acid-derivative (HADA) as a cell wall stain (39). Addition of HADA to ShyA-replete Δ6 endo cells resulted in an even distribution of staining along the cell wall (**Fig. 1B**), as expected from wild-type cell wall synthesis. In contrast, depleting ShyA resulted in a strikingly different pattern, where large patches of HADA-reactive material accumulated throughout the cell, indicative of substantial cell wall synthesis. In principle, these patches could be a remnant of incompletely-degraded cell wall material synthesized before ShyA was completely depleted, or they could reflect the activity of L,D-transpeptidases (which are able to incorporate HADA into the cell wall independent of cell wall synthesis (39, 40)). We thus repeated the staining experiment in a Δ6 endo strain lacking L,D-transpeptidases (Δ*ldtA* Δ*ldtB*). Following a 2 h depletion we added HADA for an additional hour; this still revealed an accumulation of PG patches (**Fig. 2D**), strongly suggesting that PG synthesis and incorporation continues during endopeptidase insufficiency, albeit in an aberrant, non-directional way. Quantification of PG confirmed and expanded these observations – after 2 h of ShyA depletion, cells accumulated ~18-fold more PG than ShyA-replete cells (when normalized to OD_600_) (**Fig. 1C**). Since these cells did not divide (**Fig. S2A**), PG accumulation was not correlated with an increase in cell numbers, but suggested a buildup of cell wall in the invididual, drastically enlarged ShyA-depleted cells. Consistent with a higher cell wall content, ShyA-depleted Δ6 endo cells were almost 10-fold more resistant to osmotic shock treatment (**Fig. 1D**). Thus, ShyA-depleted Δ6 endo cells not only incorporate PG, but retain higher levels of PG than the WT, possibly reflecting the lack of EP-initiated turnover processes. Similar observations have been made in autolysin-inactivated *B. subtilis*, a Gram-positive bacterium (25, 41, 42). While we cannot rule out that residual ShyA remains in the cell following depletion (at levels too low to detect above background of the non-specific band we observed via Western Blot in the same size range as ShyA, **Fig. S1**), we can at a minimum conclude that *wild-type levels* of EPs are not necessary to facilitate mass increase and incorporation of PG *per se*, but are essential for cell division and likely key for the proper, directional integration of PG into the sacculus of *V. cholerae*.

**Figure 2.**
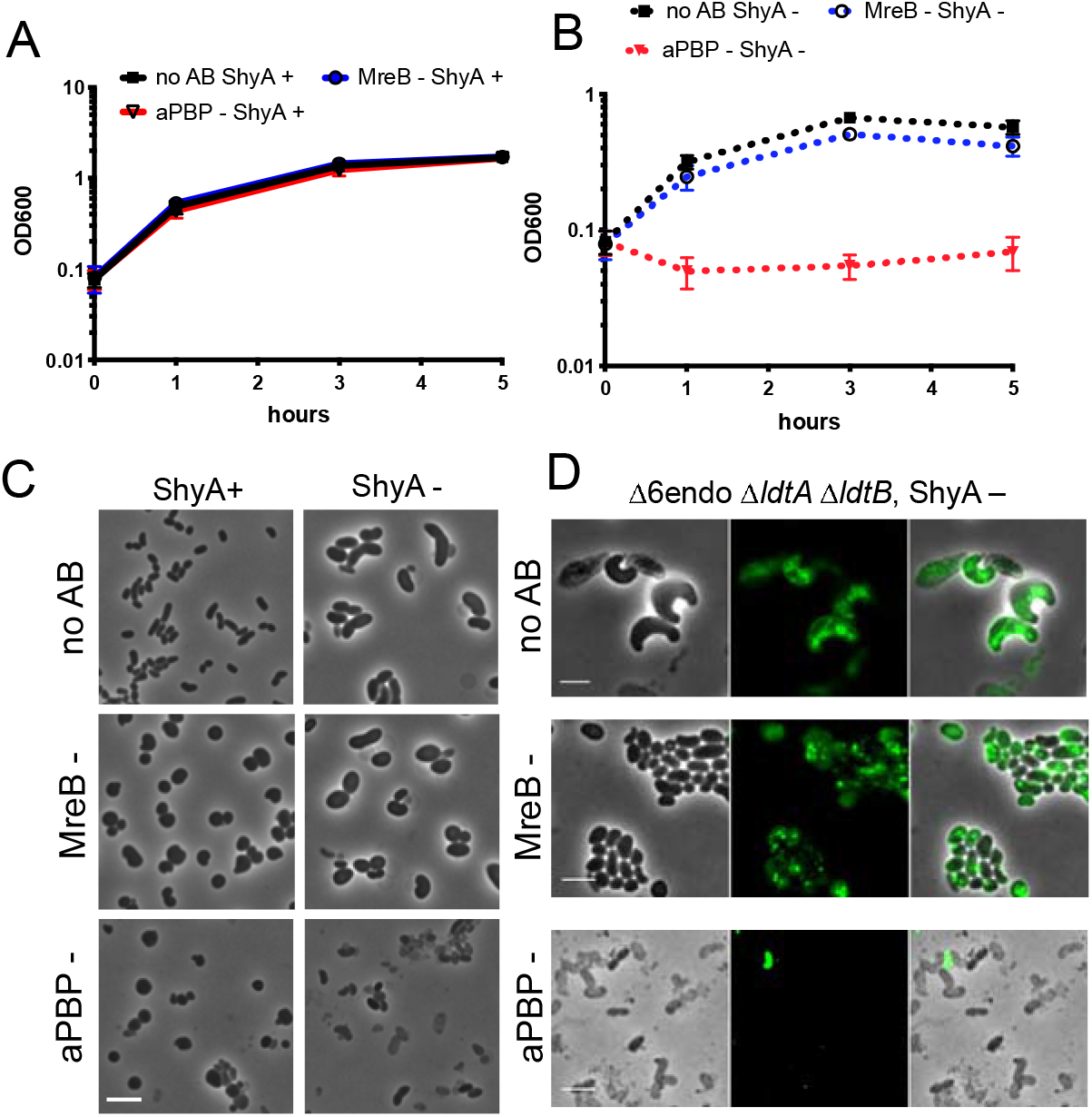
Cell mass increase and PG incorporation during EP insufficiency relies on aPBP activity. Δ6 endo grown overnight in IPTG (200 μM) was washed twice and diluted 100-fold into fresh medium containing either IPTG (ShyA+) (A) or no IPTG (ShyA -) (B) and either no antibiotic, the aPBP inhibitor moenomycin (aPBP -, 10 μg□ml^−1^, 8x MIC) or the MreB inhibitor MP265 (MreB-, 300 μM, 15 x MIC). At the indicated time points, OD_600_ was measured; after 3 h of growth, cells were also imaged (C). Data are averages of six biological replicates, error bars represent standard deviation. Scale bar, 5 μm. (D) An overnight culture of Δ6 endo Δ*ldtA* Δ*ldtB* was diluted into medium without inducer. After 2 hours of ShyA depletion, HADA (100 μM) was added for another 1 hour. Cells were then washed twice and imaged. Antibiotics MP265 (MreB -) or moenomycin (aPBP -) were added for 1 h after the 2 h initial depletion, followed by 1 h addition of HADA. Scale bar, 5 μm.

### Cell wall incorporation and mass increase in EP-deficient cells rely primarily on aPBPs

We next addressed whether EP insufficiency affected the two cell wall synthesis machines, the Rod system and the aPBPs, differentially. To this end, we repeated our Δ6 endo depletion experiment in the presence of MP265 (an inhibitor of MreB (43)) or moenomycin (an aPBP glycosyltransferase inhibitor (44)). When ShyA was expressed, mass increase proceeded at similar rates for both antibiotic treatments and the untreated control (**Fig. 2A**), while cfu/mL plateaued (moenomycin at 10 μg ml^−1^, 8 x MIC) or decreased 10-20-fold (MP265 at 200 μM, 15 x MIC) in the presence of antibiotic (**Fig. S3A**). The continued OD_600_ increase upon antibiotic exposure is consistent with our previous observations that *V. cholerae* (as well as many clinically significant Gram-negative pathogens) is remarkably tolerant to inhibitors of cell wall synthesis. Exposure to such agents causes *V. cholerae* to form cell wall-deficient spheroplasts (in the presence of aPBP inhibitors) or spheroid cells containing cell wall material (in the presence of MreB inhibitors) (30, 45). Importantly, both sphere cell types continue to increase in mass (30, 45), but fail to divide. Thus, OD_600_ continues to increase while cfu/mL stagnates or declines.

Upon ShyA depletion, mass increase in the presence of MP265 continued at a similar rate compared to ShyA-replete conditions, and we observed substantial HADA incorporation (**Fig. 2B, Fig. 2D**), suggesting that the Rod system was not absolutely required for PG synthesis during EP insufficiency. MP265-treated cells under ShyA depletion conditions qualitatively exhibited a rounder morphology than those in the untreated control, but were intact and enlarged (**Fig. 2C**). Importantly, the mass increase, HADA incorporation, and morphological aberrations were recapitulated when another Rod system inhibitor, the bPBP2 inhibitor mecillinam, was used (**Fig. S3C-F**).

In striking contrast to Rod system inhibition, aPBP inhibition via moenomycin exposure completely abrogated growth (measured by OD_600_) of ShyA-depleted Δ6 endo cells (**Fig. 2B**). This coincided with accumulation of small cells and debris (indicative of lysis), notably without strong HADA incorporation (**Fig. 2C-D**). In addition, cell viability declined rapidly in early stages (consistent with our previous observations (30)), though ultimately exhibited levels of survival similar to untreated or MP265-treated, ShyA-depleted Δ6 endo cells (**Fig. S3A-B**). In summary, our data suggest that during EP-insufficiency, the aPBPs contribute to both mass increase and sustained PG incorporation, while the Rod system is not absolutely required.

### MreB movement continues in EP-insufficient cells

The Rod-system, in conjunction with the actin homolog MreB, deposits new cell wall material during cell elongation while performing a rotational movement around the cell, apparently driven by aPBP-independent cell wall synthesis (46–48). Since the Rod system did not appear to contribute to PG synthesis during EP insufficiency, we asked whether EP depletion resulted in immobile Rod-complexes, similar to what has been observed during inhibition of cell wall synthesis (47). We constructed a functional (**Fig. S4**) mreBmsfGFP sandwich fusion in a Δ6 endo background and measured mreBmsfGFP velocity using epifluorescence and Total Internal Reflection Fluorescence (TIRF) microscopy. As a positive control, we confirmed that MP265 stopped MreB movement (**Fig. 3B**), as expected from what has been reported in *E. coli*. Mean square displacement values indicated mixed populations of diffusive MreB particles and those exhibiting directed motion under both ShyA replete and depleted conditions (**Fig. S5**). Interestingly, MreB movement continued even after 3 h of ShyA depletion (**Fig. 3D**), albeit at reduced velocity (decreasing from ~70 nm/s to ~ 40 nm/s) compared to ShyA-replete conditions (**Fig. 3C**). Our estimates of MreB velocity under ShyA replete conditions were higher than what has been reported previously for other bacteria (55 nm/s for *B. subtilis* (49) and 10 nm/s for *E. coli* (47)), perhaps reflecting species-specific differences or different properties of our sandwich fusion. Interestingly, the average size and number of MreB clusters also increased under ShyA depletion conditions (**Fig. 3E-G**), suggesting that EP depletion might affect Rod complex assembly dynamics. We conclude that similar to what has been observed in *B. subtilis* (22), EP insufficiency does not result in immediate inactivation of the Rod system, but changes its velocity and potentially the stoichiometry of its assembly.

**Figure 3.**
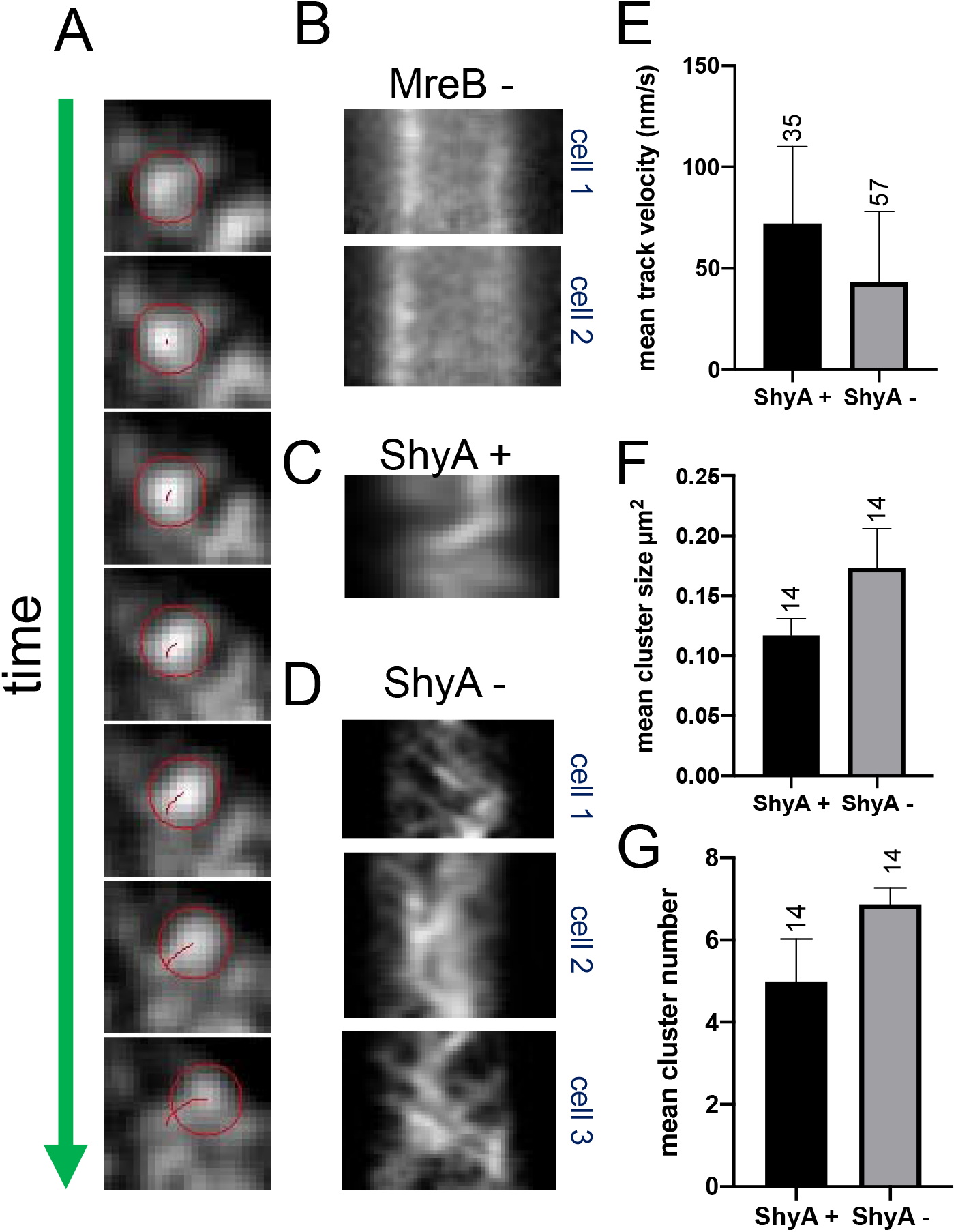
MreB movement continues during EP insufficiency. Δ6 endo **(A-D)** or Δ8 endo **(E-G)** expressing an mreBmsfGFP^sw^ fusion from its native chromosomal locus was diluted from an overnight culture grown in the presence of IPTG into growth medium without inducer (ShyA -). After 3 hours, cells were imaged using epifluorescence microscopy **(A-D)** or TIRF **(E-G)**. MreB movement was analyzed using Fiji (TrackMate). A representative single moving MreB focus track (red circle) is shown in **(A)** (frames are 2.5 s apart). Representative kymographs of MreB foci are shown for cells grown **(B)** in the presence of the MreB inhibitor MP265 (MreB -), (C) in the presence of inducer (ShyA +), or (D) in the absence of inducer (ShyA -). **(E-G)** TIRF was used to assess MreB focus velocity, mean cluster size and mean cluster number. Numbers above bar graphs represent the number of imaged cells, error bars represent standard deviation. Differences between ShyA+ and ShyA – were significant for all three graphs (t-test) at p = 0.017 (velocity), 0.021 (cluster size) and 0.028 (cluster number), respectively.

### *Complementation of EP-insufficiency in* V. cholerae *by expression of heterologous EPs*

So far, our results suggested that during EP insufficiency, aPBPs continue to synthesize PG and the Rod system may remain functional, yet does not insert substantial amounts of new PG material into the sacculus. This might suggest that the Rod system requires a physical association with one or more EPs for insertion of nascent PG material. Alternatively, EPs might catalyze PG insertion independently, *e.g*., through recognition of intrinsic PG substrate cues. To gain a better understanding of the necessity for a physical interaction, we conducted cross-species complementation experiments using an EP from *Neisseria gonorrhoeae* (henceforth “MepM_Ngo_”). This distantly related EP (a BLAST alignment indicated 29 % identity between MepM_Ngo_ and ShyA, **Fig. S6**), when heterologously expressed, is unlikely to interact with any native *V. cholerae* enzymes, and should thus allow us to isolate their EP activity from the interaction networks they might be embedded in. We expressed arabinose-inducible MepM_Ngo_ in Δ6 endo and measured differential growth in the presence of IPTG (ShyA expression) vs. arabinose (MepM_Ngo_). We found that wild-type MepM_Ngo_ was unable to rescue growth of a Δ6 endo during ShyA depletion conditions (**Fig. 4A**). However, we recently demonstrated that EPs from diverse organisms (including *E. coli* and *N. gonorrhoeae*) are produced predominantly in an inactive form due to the inhibitory function of their domain 1 and likely activated *in vivo* by an unknown mechanism (34). Heterologously expressed enzymes may not be subject to this activation pathway in *V. cholerae* (especially if the activator is a protein) and we thus instead expressed EP mutant versions with their inhibitory domain 1 deleted, rendering them constitutively active, and provided a signal sequence (ss) to ensure export to the periplasm. Surprisingly, ssMepM_Ngo_^ΔDom1^ fully complemented growth of the Δ6 endo strain to a similar degree as the native ShyA (**Fig. 4A**). Visual inspection of Δ6 cells relying on heterologous expression for growth (ara+ condition), revealed that complementation with ssMepM_Ngo_^ΔDom1^ (but not its active site mutant derivative H373A) promoted both growth (**Fig. 4A**) and the generation of rod-shaped cells (**Fig. 4B**). We sought to confirm that this apparent complementation of rod shape was still dependent on MepM_Ngo_^ΔDom1^ (rather than a mutation derepressing *shyA* in Δ6 endo). Thus, we plated all strains on agar containing IPTG, arabinose, or no inducer at the end of the experiments where we visualized cells relying on MepM_Ngo_^ΔDom1^ for growth. All strains had the same low level of spontaneous supressors able to grow in the absence of inducer (**Fig. S7**), confirming that the majority of the rodshaped cells observed when only MepM_Ngo_^ΔDom1^ was expressed are not suppressors. Interestingly, we found that heterologous expression of activated MepM from *E. coli* (ssMepM_Ngo_^ΔDom1^) was able to restore growth, but not rod-shape, to Δ6 endo *V. cholerae* (**Fig. S8**). Thus, for an unknown reason, intrinsic properties of specific EPs (or rather these mutant derivatives) define their ability to complement of Δ6 endo cell shape. In summary, these data demonstrate that heterologous expression of an activated EP can be sufficient to restore both growth and (in the case of the *N. gonorrhoeae* EP) proper cell shape to Δ6 endo cells.

**Figure 4.**
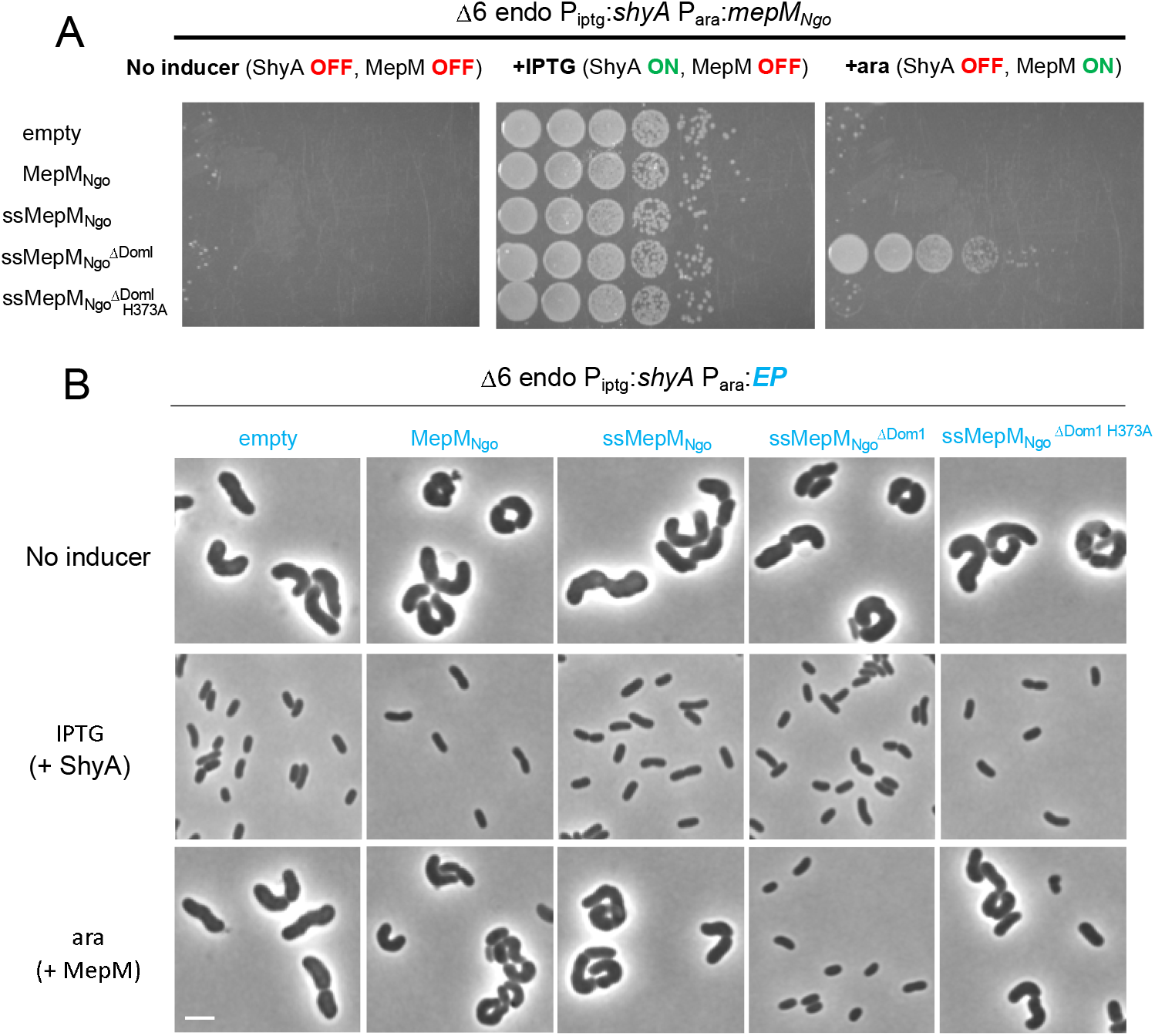
Cross-species complementation of Δ6 endo phenotypes with an EP from *Neisseria gonorrhoeae*. **(A-B)** Δ6 endo was transformed with (arabinose-inducible) pBAD33 expressing an *N. gonorrhoeae* EP (MepM_Ngo_). Derivatives of MepM_NGO_ include an N-terminal dsbA signal sequence (ss), domain 1 truncation (ΔDom1)’, or active site mutation (H373A). (**A**) Cells were washed and spot-plated on medium containing either no inducer, IPTG (200 μM) (chromsomal ShyA on), or arabinose (0.2%) (pBAD33-encoded MepM_NGO_ on). Plates were incubated at 37 °C for 24 hours and then imaged. (**B**) Cells were diluted 100-fold and grown without inducer, with IPTG (+ ShyA), or with arabinose (+MepM) for 3 hours and then imaged. Scale bar, 5 μm.

## Discussion

Bacteria must maintain a careful balance between cell wall cleavage and synthesis to promote cell elongation/division, but the exact relationship between the two cell wall synthases (Rod system vs. aPBPs) and cell wall hydrolases (*e.g*., endopeptidases) is poorly understood, at least in Gram-negative bacteria. Here, we have used EP depletion and chemical inactivation experiments to dissect the interplay between cell wall cleavage and synthesis in the cholera pathogen *V. cholerae*. Our key observation is that in *V. cholerae*, cell wall synthesis and cell expansion (but not cell division) continue upon EP depletion. This poses an apparent contradiction to data obtained in *E. coli*, where cell wall incorporation was drastically reduced after EP depletion and cells started to lyse (21). While ostensibly fundamental aspects of the coordination between cell wall synthesis and cleavage may simply not be as well-conserved as one might expect, these observations might also reflect species-specific differences in EP-independent cell wall turnover rates, and not necessarily the consequences of EP depletion *per se*. It is possible that lysis under EP-insufficient conditions in *E. coli* reflects generally higher PG degradation rates (*E. coli*, for example encodes three amidases (50), while *V. cholerae* possesses only one (51)). This would mask the underlying continued incorporation of new cell wall material in the absence of EPs. Importantly, EP depletion in *E. coli* did result in a cell volume increase prior to lysis (21), also supporting at least a transient continuation of PG synthesis during EP insufficiency in this species.

Cell wall expansion during EP-insufficiency was surprising, since presumably any form of cell wall synthesis that promotes the degree of cell expansion we observed in EP-deficient *V. cholerae* should require some form of cleavage, likely catalyzed by other autolysins. The incisions resulting from such cleavage, and/or the autolysin(s) involved appear to be of limited utility to the Rod-system, but can be exploited by the aPBPs. This suggests that aPBPs are more versatile in recognizing a variety of cell wall cuts (independent of an actual physical connection with EPs), while the Rod system primarily relies on either EP-mediated cleavage, or a physical association with EPs (see a more detailed discussion below) for substantial PG incorporation and cell expansion. These interpretations are in line with some old and several recent proposals based on data from *E. coli* that aPBPs and SEDS have separate (yet perhaps overlapping) functions during cell elongation (14, 52, 53). It has also been shown that, in *E. coli*, upregulated EP activity promotes aPBP function, likely indirectly through the creation of PG incisions that allow for an interaction between aPBPs and their OM-localized activators (28). Thus, EP cleavage may not be strictly necessary for, but can promote, aPBP activity. It is possible that under EP-insufficient conditions, the lytic transglycosylases (the other major group of cell wall cleavage enzymes that cut the polysaccharide backbone of PG (54)) create large open areas in PG that can be recognized, and patched, by aPBPs; LTG activity would be consistent with the increase in anhydro “caps” we observe in ShyA-depleted Δ6 endo PG from *V. cholerae*.

The observation (consistent with what has been shown in *B. subtilis* (22)) that MreB continues directed movement at least for some time during EP insufficiency suggests that the Rod system does not actually require wild-type EP activity for assembly and for RodA’s glycosyltransferase activity (which likely drives MreB movement). Similar to what has been proposed for *B. subtilis*, it is tempting to speculate that *V. cholerae* may use a “make-before-break” model as proposed by Höltje/Koch (35, 36) for cell elongation via the Rod system. In this model, the Rod system creates a second layer of PG that is incorporated via EPs during or after synthesis. Generation of this second layer could at first proceed independently of wild-type EP activity, but incorporation into the growing sacculus would require crosslink cleavage.

As mentioned above, our data suggest that to contribute substantially to cell expansion and PG incorporation, the Rod system requires EP activity, either through physical association or recognition of EP cut sites in PG. Our cross-species complementation experiments with an activated *N. gonorrhoeae* EP suggest that a physical association might not be strictly necessary, unless the heterologously expressed (and truncated) enzyme does somehow directly interact with the *V. cholerae* Rod system. We thus consider a model plausible where rather than (or in addition to) co-ordinating with cell wall synthases directly, EPs can somehow specifically recognize and preferentially cleave old PG that is adjacent to nascent PG. Though highly speculative, our observation that the corresponding *E. coli* homolog does not complement cell shape might reflect different levels of activity – since EPs can promote aPBP activation (at least in *E. coli*) (28), overexpression of a more active EP might divert PG precursor flux away from the Rod system towards aPBPs to a higher degree than the *N. gonorrhoeae* enzyme, incapacitating a cell’s ability to elaborate a rod shape.

An important caveat to the complementation experiments, is that the Δ6 endo strain still maintains a copy of *shyA* under IPTG control. While the lac promoter is tightly repressed in the absence of inducer, a small number of molecules under its control might still be produced (55). ShyA is produced predominantly as an inactive precursor and the signal for activation is unkown (34). It is conceivable that complementation with a heterologously expressed EP might somehow enhance activation of this leaky background of ShyA molecules*, e.g*. if there is a positive feedback loop between cell wall cleavage and native EP activation.

Taken together, our data suggest that two main cell wall synthases, the aPBPs and the Rod system have differential relationships with autolysins, and especially endopeptidases. As such, our data provide additional support for the emerging theme of at least partially differential roles of the aPBPs and the Rod system during cell elongation.

## Acknowledgements

This work was supported by the National Institutes of Health/NIGMS through R01 GM130971 to TD. Research in the Cava lab is supported by MIMS, the Knut and Alice Wallenberg Foundation (KAW), the Swedish Research Council and the Kempe Foundation. Research in the VanNieuwenhze lab is supported by the NIH through R01 GM113172 and R35 GM136365. TIRF imaging was supported by grant NSF 1428922 to the BRC imaging facility at Cornell.

## Materials and Methods

### Bacterial growth conditions

Cells were grown by shaking (200 rpm) at 37°C in 5 mL of LB in borosilicate glass tubes (14 mL capacity) unless otherwise indicated. Where appropriate, antibiotics were used as the following concentrations: streptomycin, 200□μg□ml^−1^; ampicillin, 100□μg□ml^−1^; chloramphenicol, 5□μg□ml^−1^; moenomycin, 10 μg ml^−1^; MP265, 300 μM; and mecillinam, 10 μg□mΓ^−1^. IPTG (200□μM) and arabinose (0.2%) were added for induction of P_iptg_ and P_ara_ promoters, respectively.

### Plasmid and strain construction

All bacterial strains and oligonucleotides used in this study are summarized in **Table S1**. All *Vibrio cholerae* strains are derivatives of El Tor strains N16961 (56) or E7946 (57), the latter was used for chitin-induced transformation.

Δ6 endo construction is reported elsewhere (30). Other strains were constructed by chitin-induced transformation of linear PCR products as described in (58). A chloramphenicol (*chl*) resistance cassette insertion into the gene *vc1807* (a well-established neutral locus) was used as the primary selector. The transforming fragment for vc1807::chl was constructed by amplifying upstream and downstream homology regions using primers PD079/PD097 and PD098/PD082, respectively. The *chl* gene coding for chloramphenicol acetyl transferase was amplified from pBAD33 (59) with primers PD095/PD096 and fused with the flanking homologies of vc1807 via isothermal assembly. For antibiotic resistance gene swapping, a vc1807::trim allele was also produced by amplifying upstream (using primers TDP597/598) and downstream (primers TDP601/602) homologies of vc1807 and fusing them with a trimR cassette amplified from *V. cholerae* Haiti (60) (primers TDP599/600) using SOE PCR with primers TDP603/604.

To construct a functional MreB-msfGFP-MreB sandwich fusion, upstream (primers PD056/PD074) and downstream (primers PD071/PD057) homologies were amplified from the *V. choerae* genome and fused via isothermal assembly with msfGFP (amplified with primers PD054/PD055). Analogous to a published *E. coli* MreB-msfGFP sandwich fusion (47), we replaced glycine 228 of MreB with this msfGFP. To enhance the probability of success of finding a functional fusion, we used semi-degenerate primers to generate a library of possible linker sequences. Flanking homologies, MreB and msfGFP were first fused using isothermal assembly (61) and then amplified using nesting primers PD104/PD105. The resulting upstream-MreB-linker-msfGFP-linker-MreB-downstream PCR fragments were transformed into E7946 using chitin transformation with vc1807::chl as the primary selector. 96 colonies were tested for growth rate and clone M2C was chosen for further experiment due to its wild-type growth behavior. The linkers of this fusion construct were sequenced (coding for DGVGG upstream of msfGFP and GTPIP downstream).

Δ8 was constructed by transforming endopeptidase deletion PCR products into a parental mreB::mreBmsfGFP ΔlacZ::P_IPTG_:*shyA* strain. Deletion scars were amplified from Δ6 endo and introduced in two steps into this parental background via chitin transformation. The following primers were used to amplify the EP deletion fragments: *shyA* (TDP577/578), *shyC* (TDP581/582), *shyB* (TDP579/580), *vc1537* (TDP583/584), *vc0843* (tagE1) (TDP587/588), *vca1043* (tagE2) (TDP585/586). PBP4 and PBP7 deletions were introduced into Δ6 endo by amplifying PCR fragments with upstream and downstream homologies fused by a linker for chitin-mediated transformation. PBP4 upstream homology (TDP680/TDP681) and downstream homology (TDP682/TDP683) were fused using SOE PCR with nesting primers (TDP691/TDP692). PBP7 upstream homology (TDP676/TDP677) and downstream homology (TDP678/TDP679) were fused using SOE PCR with nesting primers (TDP693/TDP694). For the Δ*ldtA* Δ*ldtB* strain, deletion scars were amplified from strain FC670 (40) using primers TDP654/55 (*vc1268*) and TDP656/57 (*vca0058*), respectively, and transformed into Δ6 endo using chitin transformation as described above.

Δ8 exhibited a very low transformation efficiency and we thus introduced the Δ*vc1269* deletion using homologous recombination with a suicide plasmid pCVD442 as described (62). In brief, upstream and downstream homologies of vc1269 were amplified using primers TD810/TD811 and TD812/TD813. These fragments were cloned into Xba1-digested pCVD442 using isothermal assembly. pCVD442(Δvc1269) was then introduced into Δ8 endo via biparental mating (using SM10 as a donor strain) by mixing 10 μL of each donor and recipient, followed by 6 h incubation at 37 °C, followed by selection for single crossover strains and against the donor strain by plating on LB plates containing carbenicillin (100 μg□ml^−1^), streptomycin (200 μg□ml^−1^) and IPTG (200 μM). A single colony from the first crossover plate was then picked and streaked out on a plate containing sucrose (10 %), streptomycin (200□μg ml^−1^) and IPTG (200 μM). This plate was incubated at ambient temperature for 3 days, after which 16 colonies were tested for the correct knockout construct using vc1269 flanking primers TD814/TD815.

All plasmids were built using isothermal assembly (61). Genes were cloned into pBADmob (a mobile pBAD33 derivative) using the following primer pairs: MepM_Ngo_, TDP1365/TDP1367, ssMepMNgo, SM861/SM862; ssMepMNgo^Δdom1^, SM859/SM860; MepM_ECO_, TDP1342/TDP1340; and MepM_ECO_^Δdom1^, TDP1341/TDP1340. The H373A point mutation was introduced into pBAD plasmids carrying NGO1686 derivatives via Q5 site-directed mutagenesis (NEB, Ipswitch, MA, cat #E0554S) with primer pair TDP1652/TDP1653. Plasmids were conjugated into *V. cholerae* using donor *E. coli* strains (SM10 lamda pir or MFD lamda pir).

### Phase contrast microscopy and HADA staining

Cells were harvested (2 min at 12,000 rpm), spotted on a 0.8% agarose pad containing PBS and imaged on a Leica Dmi8 inverted microscope. For HADA experiments, Δ6 endo cells were grown in the presence of 50 μM HADA (3-[[(7-Hydroxy-2-oxo-2*H*-1-benzopyran-3-yl)carbonyl]amino]-D-alanine hydrocholoride), washed once by pelleting cells (2 min at 12,000 rpm) and resuspending in fresh LB. HADA stain was imaged in the DAPI channel (395 nm [excitation]/460 [emission]) at 1 s exposure.

### Endopeptidase depletion experiments

EP depletion strains were grown overnight in LB broth containing 200 μM IPTG. The next day, cells were washed 2x by pelleting (2 min at 12,000 rpm) and resuspending in LB broth without inducer. Cells were then diluted 100-fold into fresh LB containing either 200 μM IPTG (ShyA +) or no inducer (ShyA -). Where indicated, antibiotics were used at 10 μg□m^−1^ (moenomycin, mecillinam) or 200 μM (MP265).

### Single particle tracking by TIRF imaging

The Δ8 endo mreB::mreBmsfGFP^sw^ strain with chromosomally expressed MreB-msfGFP was grown shaking at 37°C in LB medium supplemented with 100μM IPTG overnight. The saturated cells were diluted (1:100) into fresh LB in two groups (with 100μM IPTG for ShyA expression or without IPTG for ShyA depletion). After 2 hours of shaking (220 rpm) incubation at 37°C, cells were harvested and spotted on a 0.8% agarose pad containing M9 medium. Time-lapse TIRF imaging was performed on a Zeiss Elyra equipped with an inverted Axio Observer.Z1 microscope and a 100x 1.46 oil objective. The objective was heated at 37°C during image acquisition. The exposure time was 100 ms and inter-frame intervals were 2 s over a 2-min recording. The movement of MreB-msfGFP was analyzed using single particle tracking software ImageJ TrackMate (63) and MATLAB msdanalyzer (64).

The mean square displacements (MSD) of particle trajectories were calculated using the msdanalyzer package and the motion types were analyzed through log-log fitting (64). By setting the R^2^ coefficient > 0.8, individual MSD curves were fitted and the values of anomalous diffusion coefficient (α) indicates that MreB particles exhibit a mix of dynamic behaviors (confined diffusion, 0.1 ≤ α < 0.9; simple diffusion, 0.9 ≤ α < 1.1; directed motion, α ≥ 1.1) (65).

### Peptidoglycan analysis

PG samples were analyzed as described previously (66). Briefly, 50 mL cultures of Δ6 endo were grown to early/mid exponential phase with or without IPTG (200 μM) for 2h, harvested and boiled in 5% SDS for 1 h. Sacculi were repeatedly washed by ultracentrifugation (110,000 rpm, 10 min, 20°C) with MilliQ water until SDS was totally removed. Samples were treated with 20 μg Proteinase K (1 h, 37 °C) for Braun’s lipoprotein removal, and finally treated with muramidase (100 μg ml^−1^) for 16 hours at 37 °C. Muramidase digestion was stopped by boiling and coagulated proteins were removed by centrifugation (14,000 rpm, 10 min). For sample reduction, the pH of the supernatants was adjusted to pH 8.5-9.0 with sodium borate buffer and sodium borohydride was added to a final concentration of 10 mg□ml^−1^. After incubating for 30 min at room temperature, the samples pH was adjusted to pH 3.5 with orthophosphoric acid.

UPLC analyses of muropeptides were performed on a Waters UPLC system (Waters Corporation, USA) equipped with an ACQUITY UPLC BEH C18 Column, 130Å, 1.7 μm, 2.1 mm X 150 mm (Waters, USA) and a dual wavelength absorbance detector. Elution of muropeptides was detected at 204 nm. Muropeptides were separated at 45°C using a linear gradient from buffer A (formic acid 0.1% in water) to buffer B (formic acid 0.1% in acetonitrile) in an 18-minute run, with a 0.25 ml/min flow.

Relative total PG amount was calculated by comparison of the total intensities of the chromatograms (total area) from three biological replicas normalized to the same OD600 and extracted with the same volumes. Muropeptide identity was confirmed by MS/MS analysis, using a Xevo G2-XS QTof system (Waters Corporation, USA). Quantification of muropeptides was based on their relative abundances (relative area of the corresponding peak) normalized to their molar ratio.

### Western Blotting

Whole cell lysates (15 μg) were resolved by 10% SDS-PAGE and the proteins were transferred to a PVDF membrane using a semi-dry transfer system (iBlot 2, Invitrogen). The membrane was then blocked overnight with blocking solution containing 4% milk (dry milk dissolved in 20 mM Tris-HCl (pH 7.8), 150 mM NaCl, 0.1% Triton X-100). Next day, the membrane was incubated with anti-ShyA polyclonal antibody (1: 5,000, produced by Pocono Rabbit Farm & Laboratory, PA) for two hours and then washed twice with 1xTBST (20 mM Tris-HCl (pH7.8), 150 mM NaCl, 0.1% Triton X-100). The washed membranes were then incubated with anti-rabbit secondary antibody (1:15,000, Li-Cor cat# 926-32211) for 1 hour. Membranes were then washed three times with 1xTBST, scanned on an Odyssey CLx imaging device (LI-COR Biosciences) and visualized using Image Studio™ Lite Ver 5.2 software (Li-Cor) for signal quantification.

---

**Figure S1.**
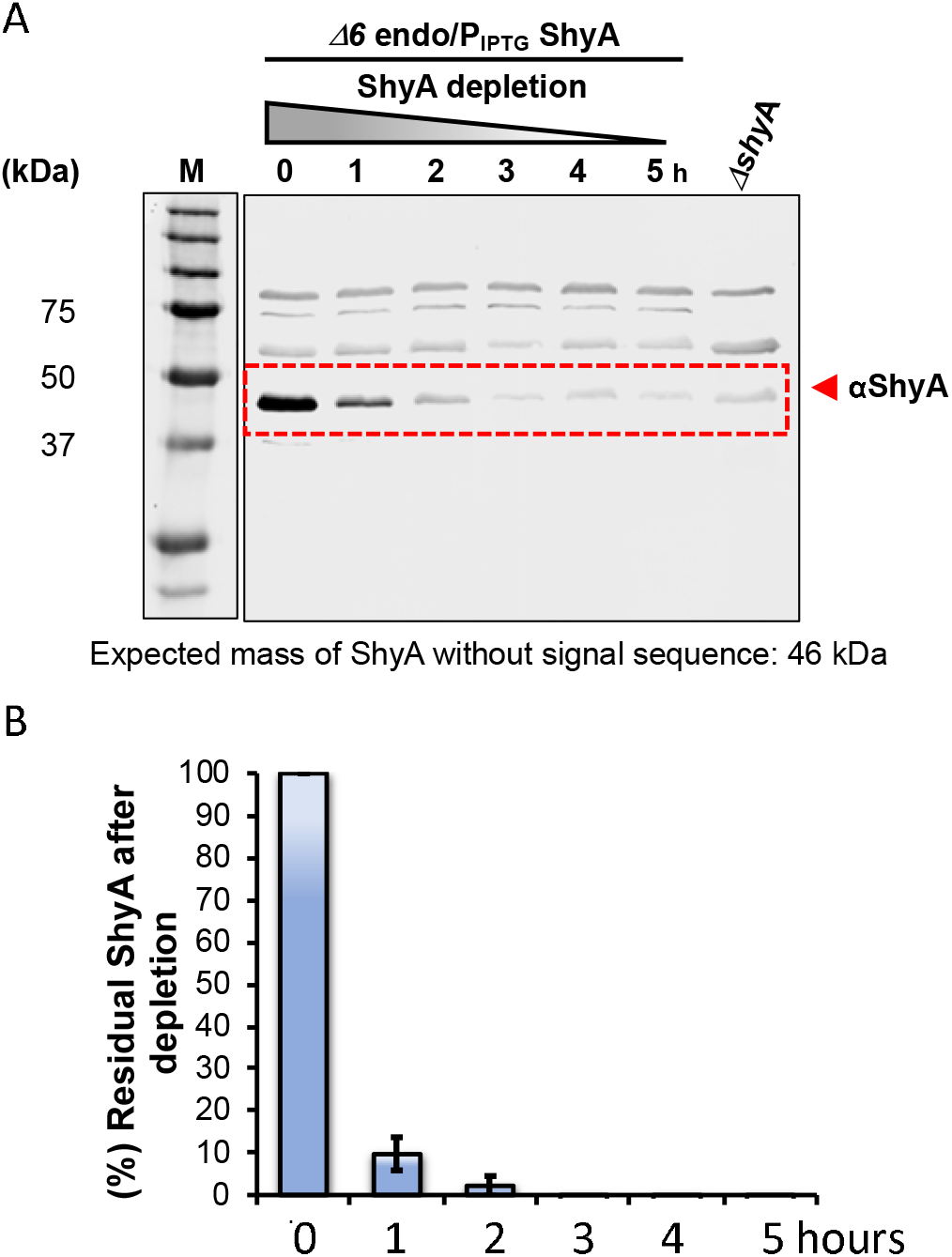
ShyA levels after depletion in Δ6 endopeptidase mutant. **(A)** Wild-type, Δ6 endo, and Δ*shyA* strains were grown overnight in LB broth containing 200 μM IPTG, then washed three times with fresh LB. Cells were then diluted 100-fold into 150 ml pre-warmed LB medium without IPTG for ShyA depletion. Samples were collected at indicated time points. For Western Blotting, cell extracts (adjusted to same protein concentration) were separated on 10% SDS-PAGE gels and subjected to Western Blot analysis using ShyA polyclonal antibody (B) ShyA band intensities were quantified (ImageJ) and the intensity value of the non-specific background band detected in the Δ*shyA* mutant was subtracted. Residual ShyA protein levels were normalized to non-depleted ShyA at 0 h (100%).

**Figure S2.**
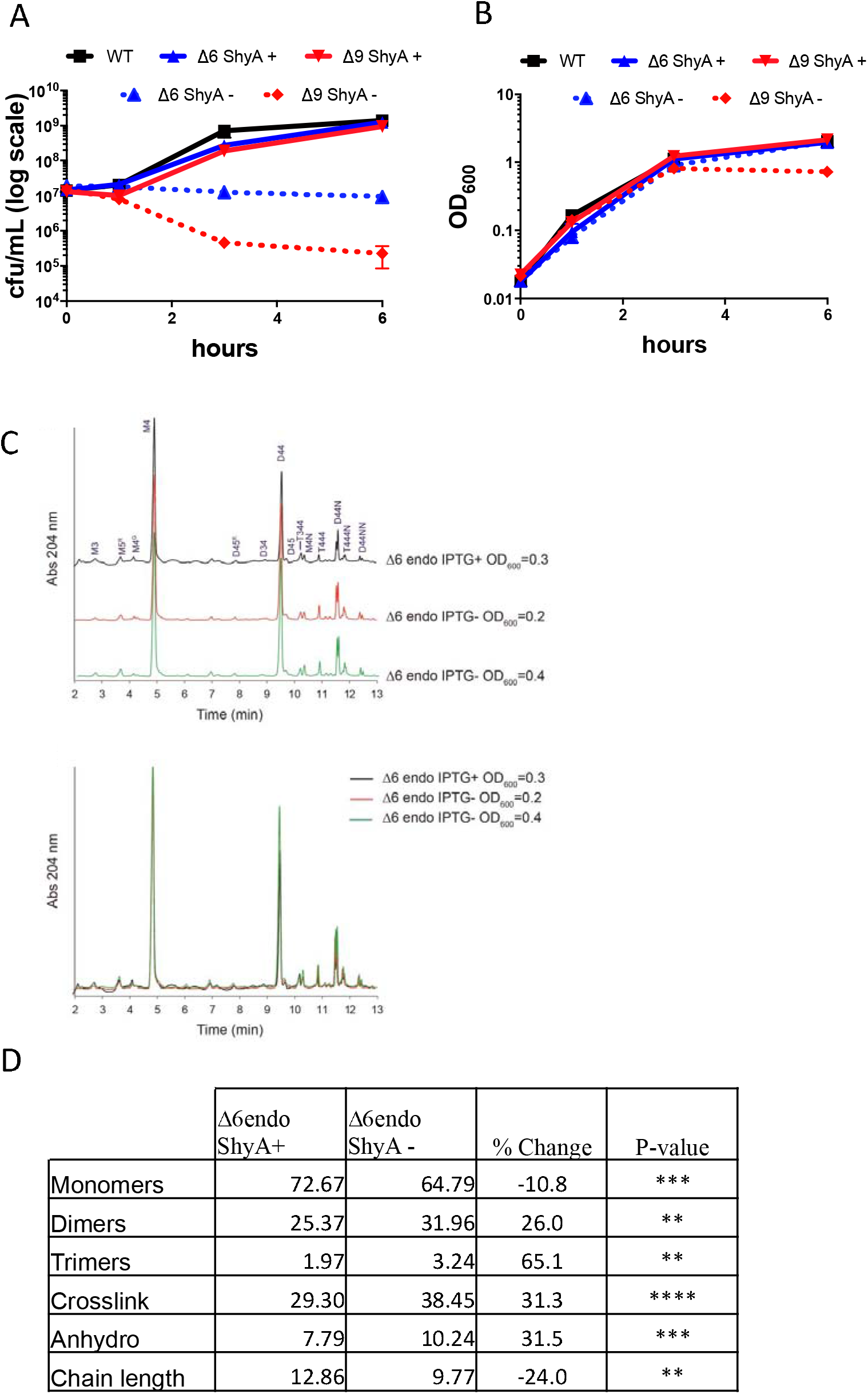
Effects of EP depletion on growth, survival and PG composition. N16961 Δ6 endo or E7946 Δ9 endo (mreB::mreBmsfGFP^sw^) was grown overnight in IPTG (200 μM), washed twice, and diluted 100-fold into fresh medium with (ShyA +) or without (ShyA -) inducer. At the indicated time points, OD_600_ (**A**) was measured via spectrophotometry and cells were diluted serially onto LB IPTG (200 μM) plates to determine colony forming units per mL (**B**). Data are averages of 3 biological replicates, error bars represent standard deviation. (**C)** Chromatogram of PG composition of N16961 Δ6 endo cells harvested after 2 hours with (ShyA +) or without (ShyA -) IPTG (200 μM). **(D)** The table summarizes the relative molar abundance (%) of monomers, dimers, trimers shown in the chromatogram. Data regarding the % of crosslinkage (proportion of crosslinked peptide side chains, calculated on dimers and trimers content) is also included. Anhydro muropeptides (with a residue of (1-6 anhydro) N-acetyl muramic acid) are the terminal subunits of the sugar chains and hence used to calculate the chain length. Values are mean of three biological replicates. Percent change was calculated relative to the IPTG-treated sample and p-values were generated using a multiple comparisons t-test (****, p < 0.0001; ***, p < 0.001, ** p < 0.01, * < 0.05).

**Figure S3.**
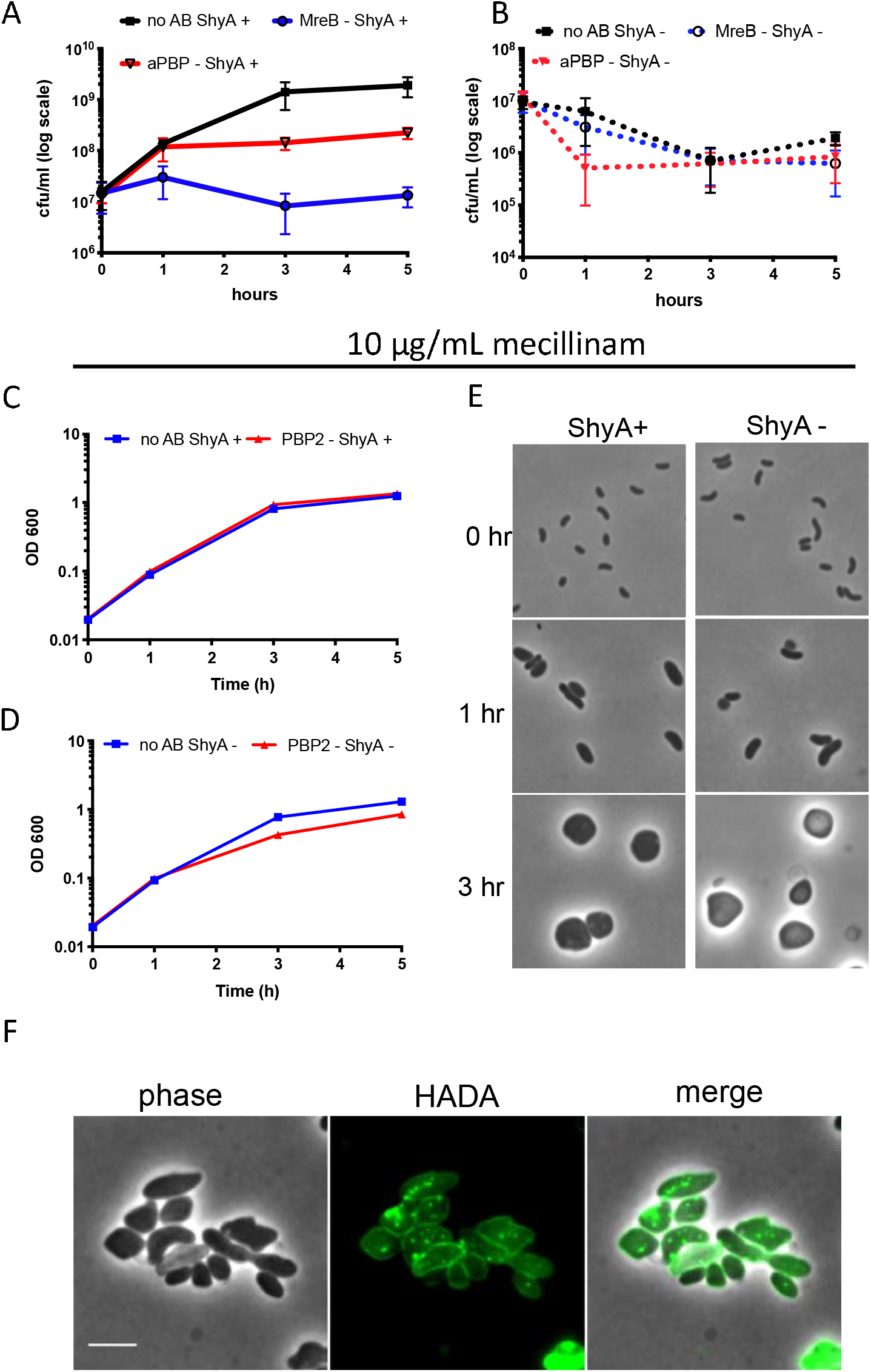
Mass increase during EP insufficiency relies on aPBPs, but not the Rod system. Δ6 endo was grown overnight in IPTG (200 μM), washed twice, and diluted 100-fold into fresh medium containing either IPTG (ShyA+) **(A)** or no IPTG (ShyA -) **(B)** and either no antibiotic, the aPBP inhibitor moenomycin (aPBP -, 10 μg□ml^−1^, 8x MIC) or the MreB inhibitor MP265 (MreB -, 300 μM, 15 x MIC). At the indicated time points, cells were diluted serially and spotted onto LB plates containing IPTG (200 μM). Data are averages of six biological replicates, error bars represent standard deviation. **(C-E)** In a similar experiment, Δ6 endo was treated with mecillinam (10 μg□ml^−1^, 20x MIC). At the indicated time points, OD_600_ **(C-D)** was measured via spectrophotometry and cells were harvested and spotted on a 0.8% agarose pad containing PBS for phase contrast microscopy **(E)**. **(F)** The cell wall was stained using HADA as described for **Fig. 2D**. Scale bar, 5 μm.

**Figure S4.**
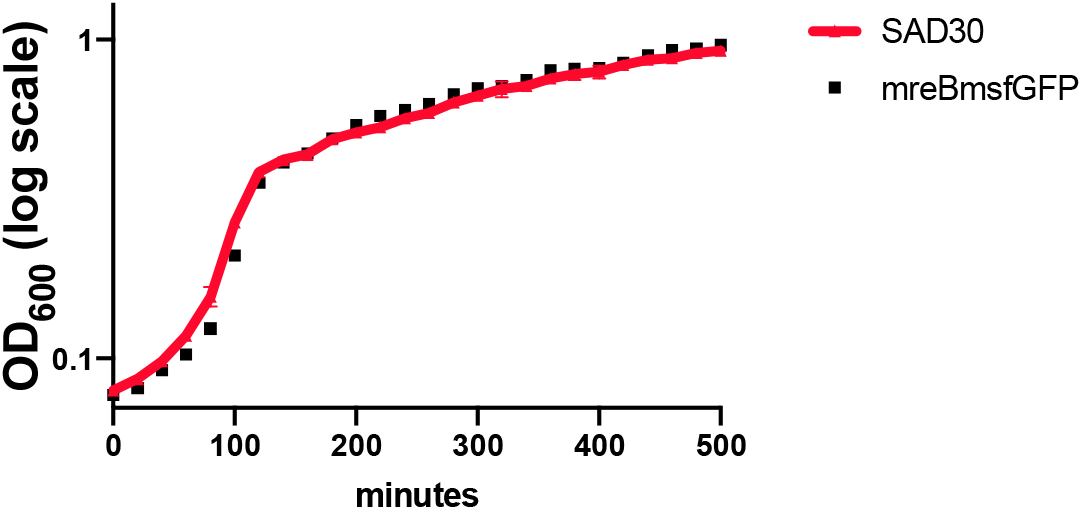
Growth of the mreBmsfGFP strain compared to wild-type. Wild-type E7946 and mreBmsfGFP-containing derivative were grown overnight in LB. Cells were diluted 1000-fold into fresh medium and 200-μl of each was loaded into a 100-well plate. Growth of each culture was monitored by optical density at 600 nm (OD_600_) in a Bioscreen C plate reader (Growth Curves America).

**Figure S5.**
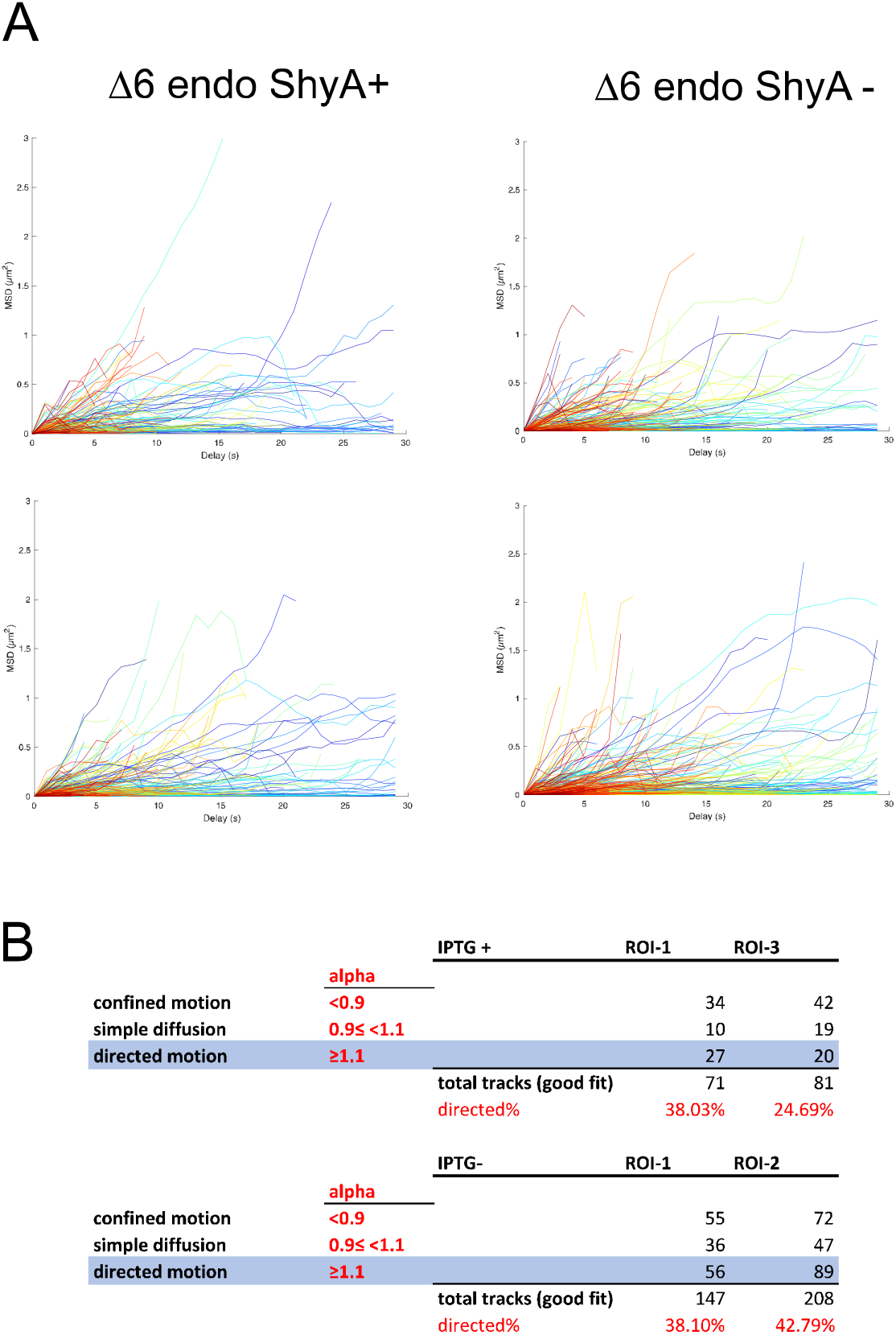
Mean square displacement analysis. Δ6 endo mreBmsfGFP^sw^ was grown with (ShyA +) or without (ShyA -) IPTG and imaged using TIRF microscopy. **(A)** Example MSD curves for 2 regions of interest (ROIs) for each, the IPTG+ and IPTG condition, **(B)** alpha values and % of MreBmsfGFP patches exhibiting directed motion.

**Figure S6.**
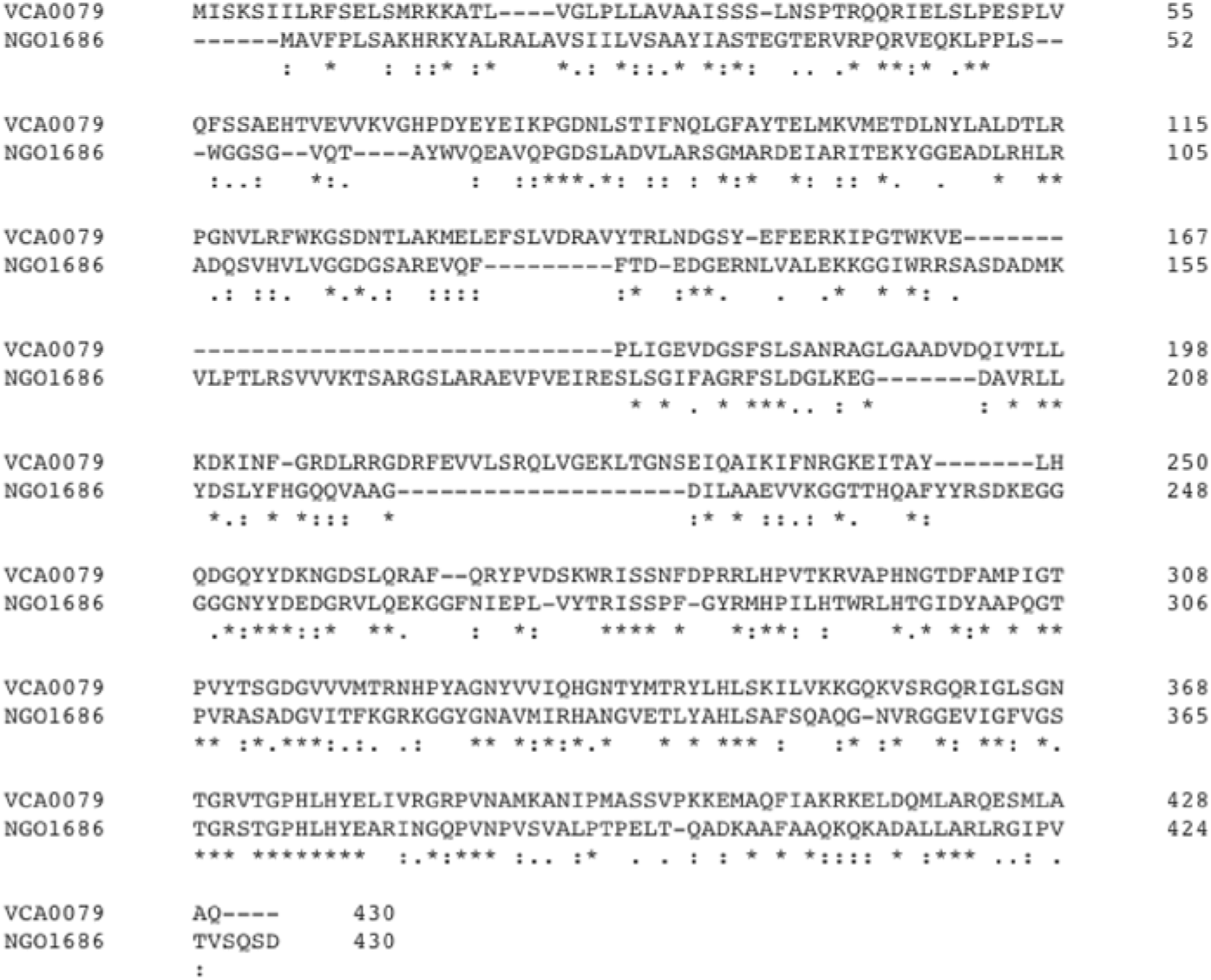
ShyA and MepMNGO amino acid sequence alignment. Amino acid sequences of *V. cholerae* endopeptidase ShyA (VCA0079) and *N. gonorrhoeae* ortholog MepM (NGO1686) were aligned in Clustal Omega (67). Numbers indicate the amino acid position relative to the start codon and alignment gaps are denoted with dashes. Symbols indicate the similarity of aligned residues: identical (*), strong similarity (:), and weak similarity (.).

**Figure S7.**
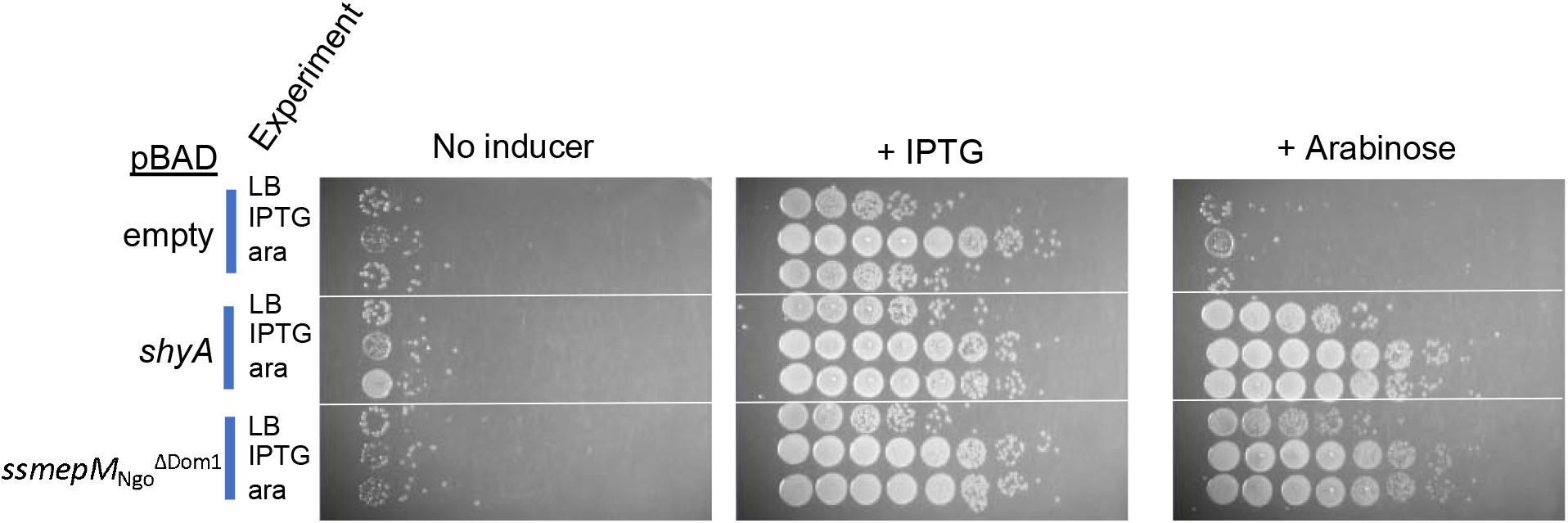
Low suppressor background in Δ6 endo cells. Δ6 endo carrying pBAD33 (arabinose-inducible) expressing the indicated contructs (“pBAD” column) were diluted into fresh medium containing either no inducer, IPTG or arabinose (ara) and grown for 3 hours (“experiment” column). Cells were then spot-plated on medium containing either IPTG (200 μM, ShyA expressed), arabinose (0.2 %, MepM_Ngo_ expressed) or no inducer. Plates were incubated at 37 °C for 24 hours and then imaged.

**Figure S8.**
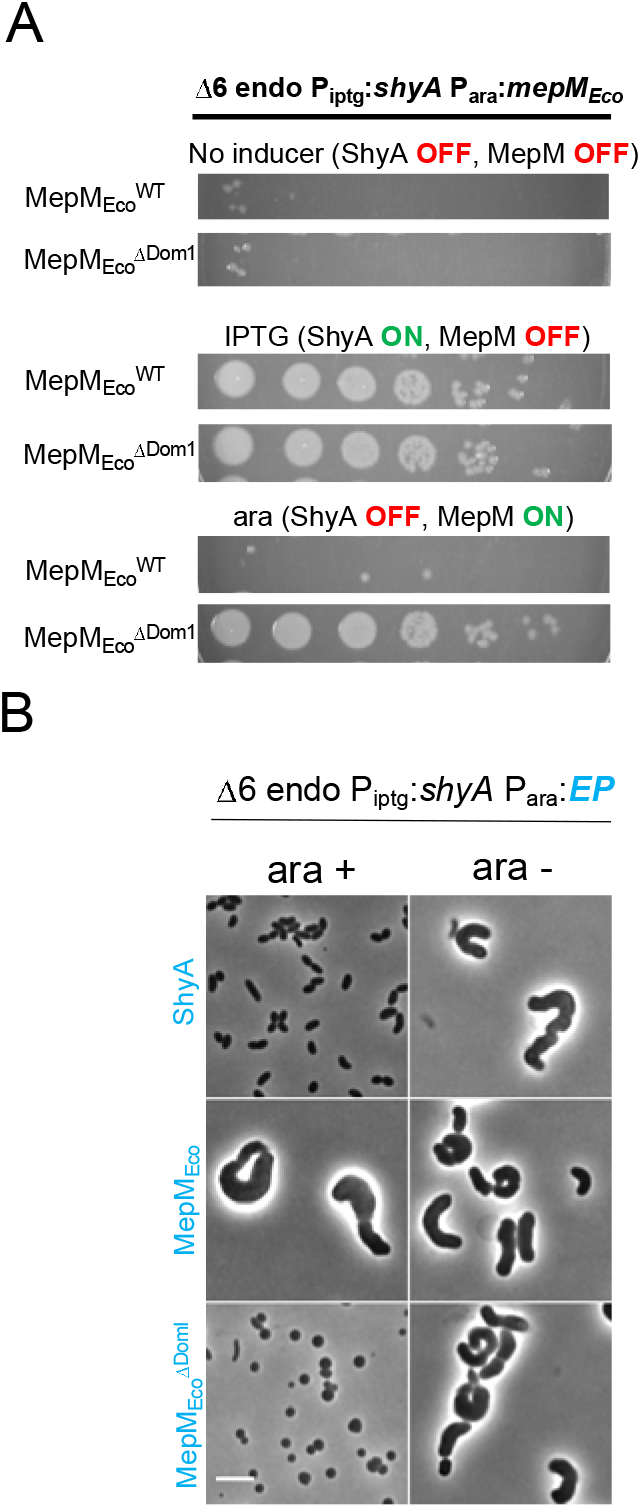
Cross-species complementation of Δ6 endo phenotypes with an EP from *Escherichia coli*. **(A)** Δ6 endo carrying pBAD33 (arabinose-inducible) encoding MepM_Eco_ or its Δdomain 1 derivative, was diluted and spot-plated on medium containing either IPTG (200 μM, ShyA expressed), arabinose (0.2 %, heterologous EP expressed) or no inducer. Plates were incubated at 37 °C for 24 hours and then imaged. (**B**) Δ6 endo carrying the indicated EP under control of an arabinose-inducible promoter was grown without IPTG (chromosomal ShyA off) and with arabinose (pBAD33-encoded EP on) for 3 hours and then imaged. Scale bar, 5 μm.

